# Transcriptomic Profiling of Old Age Sarcoma Patients using TCGA RNA-seq data

**DOI:** 10.1101/2025.01.03.631189

**Authors:** Vidhyavathy Nagarajan, Shreya S. Karandikar, Mary S.J. Dhevanayagam

## Abstract

Sarcoma is a rare malignancy with poor prognosis, especially in older patients (≥ 65 years) as seen in our preliminary analysis and some previous studies. Moreover, these patients have limited treatment options due to therapy-associated adverse effects and altered tumor micro-environment, which could be associated with their lower prognosis. Studying the underlying biology that drives cancer progression in these patients will help design personalized therapy and improve outcomes for them. This study aims to analyze TCGA-SARC RNA-seq data for characterizing the transcriptomic profile of older age (OA: ≥ 65 years) compared to younger age (YA: 18-65 years) sarcoma patients. RNA-seq and clinical data of sarcoma patients were acquired from TCGA, and the samples were grouped as OA (≥ 65 years) and YA (18-65 years) patients. Differential gene expression analysis, pathway analysis, transcription factor enrichment analysis, gene-specific survival analysis and network analysis were performed. When comparing the gene expression profiles of the 108 OA and 154 YA patients, significant differentially regulated genes (n=733), transcription factors (n=10), hub genes (n=10) and the pathways that characterize the former were identified. Furthermore, 16 dysregulated genes were found that were significantly associated with a poor prognosis in OA sarcoma patients. In accordance with existing evidence of an altered tumor microenvironment in older-age cancer patients, the identified significant genes are associated with the regulation of certain important tumorigenic pathways such as EMT (epithelial-to-mesenchymal transition), calcium signaling, angiogenesis, ECM (extracellular matrix) degradation, Wnt/*β*-catenin pathways, suggesting the potential cause for lower prognosis in the OA patients. Thus, these findings pave the way to characterize the OA sarcoma patients which can be validated by multi-omics analysis and clinical studies in the future, in turn providing improved treatment options and survival for the same.

## 1 Introduction

Sarcomas are neoplasms, accounting for 1% of adult cancers and 15% of pediatric cancers (18 years), that originate in the mesenchyme including bone, cartilage, muscle, and connective tissues of the body [1, 2]. They are classified into more than 100 subtypes, with 80% classified as soft tissue sarcoma (STS) and 20% as bone sarcomas (BS) [1, 3, 4]. Soft-tissue sarcomas (STS) are a heterogeneous group of mesenchymal malignancies that primarily attack the retroperitoneum, extremities, trunk, and head and neck attributing significant therapeutic challenges [5]. Bone sarcomas (BS) are very rare, only 0.2% of all malignant tumors found, and arise in mesenchyme cells of the body’s skeletal system [6].

Due to the high heterogeneity in the subtypes, very few studies have conducted an overall survival (OS) analysis for all sarcomas, but many of them examined and reported the survival of STS and BS separately [7, 8]. The net 5 year OS of STS and BS patients were reported as 53% and 61% respectively excluding gastrointestinal sarcomas (which was higher at 70% due to development targeted therapies) [7]. Other studies showed that BS patients had an average survival rate ranging from 56% to 70% [9–11], while STS patients had a survival rate ranging from 56% to 79% [12–14].

Although there has been some increase in the survival rate of sarcoma patients over time, it depends on multiple prognostic factors and has reached a stagnant rate for some sarcomas [7]. Apart from the tumor site, which varies very widely, age of incidence is an important prognostic factor involved in determining survival rate for both STS and BS patients [6, 8, 13, 15–17]. Studies show that the incidence rate of sarcoma depends on the histologic subtype and is more common among children (*>* 18 years) and elderly people (*>* 65 years) [17–20]). Despite the high occurrence of sarcomas in the older age (OA) patients, their survival rate is poor compared to younger patients [7, 8, 13, 21–25]. There have been a few contradictory results as well reported previously supporting a lower survival rate in younger patients [26–30].

In general, sarcomas are difficult to manage and treat due to misdiagnosis (difficult to differentiate from other cancer types), late diagnosis (absence of early symptoms), high heterogeneity, aggressiveness and resistance to current treatments [31]. Currently, surgery combined with pre- or post-operative therapy is the most promising therapy for localized sarcoma [32–34]. Radiation and chemotherapy are ineffective for metastatic sarcomas, which affect one-third of patients and account for 20% of recurrence rates [35]. Further, there is a under-representation of elderly patients in clinical trials and less aggressive therapies are used to prevent adverse effects [18, 36–39]. Additionally, OA STS patients are initially presented with higher grade and larger tumors making it difficult to treat them [18, 40–42].

Studies have shown that age is a prognostic marker in colorectal cancer (CRC), liver and pancreatic cancer as well [43–45]. Though undertreatment of OA patients impacts survival, other factors like the tumor biology of older patients also plays a role in management of cancer. Aging is associated with immunosenescence (decrease in anti-tumor cell mediated immunity), alterations in ECM (extracellulat matrix), increased inflammation, all of which, promote tumorigenesis process [15, 46, 47]. Further, it was reported that the tumor microenvironment in the elderly promotes metastasis pathways such as remodelled ECM along with immunosenescence [48–50]. It is evident that there is a change in molecular level that favors tumorigenesis in older patients. In this study we aim to study the changes in transcriptomic level and identify significant genes undergoing changes in their expression.

## 2 Background

Due to the collection of conflicting evidence regarding the prognosis of OA and YA sarcoma patients, we first sort to understand the prognostic difference based on age using TCGA-SARC data (clinical data from TCGA database) [7, 8, 13, 21–30]. The subtypes of sarcoma included in the dataset mainly includes, STS (leiomyosarcoma, synovial sarcoma, liposarcoma, fibromyxosarcoma, malignant peripheral nerve sheath tumor (MPNT), fibromatosis, malignant fibrous histiocytoma), undifferentiated sarcoma, and giant cell sarcoma which is illustrated in the Supplementary Figure 1. It is seen that patients above the age of 65 (OA) show lower prognosis compared to patients between 18-65 years (YA) based on Cox regression analysis (p=0.0000938, HR - 2.22, 95% CI - [1.488, 3.316]) and Kaplan-Meier (KM) log rank test analysis (p=0.00006). Further, the median survival for OA is about 3 years while that for YA is 7 years. The KM analysis plot as illustrated in (Fig. 2.0.1) shows that the OA patients have a lower survival probability compared to YA.

Merry *et al.,* 2021 [51] summarizes the potential of transcriptomic biomarkers in sarcoma patient stratification and treatment. Few studies have identified biomarkers based on either transcriptomic analysis alone or integrated with other analyses, such as proteomics, mutational analysis, etc., to study the patient prognosis [52–57]. However, transcriptomic biomarkers that distinguish the OA patients from YA have not yet been characterised. Hence, in this study we aim to identify the transcriptomic biomarkers and significant pathways that are disrupted in the OA patients and associated with their lower prognosis.

**Fig 2.1.**
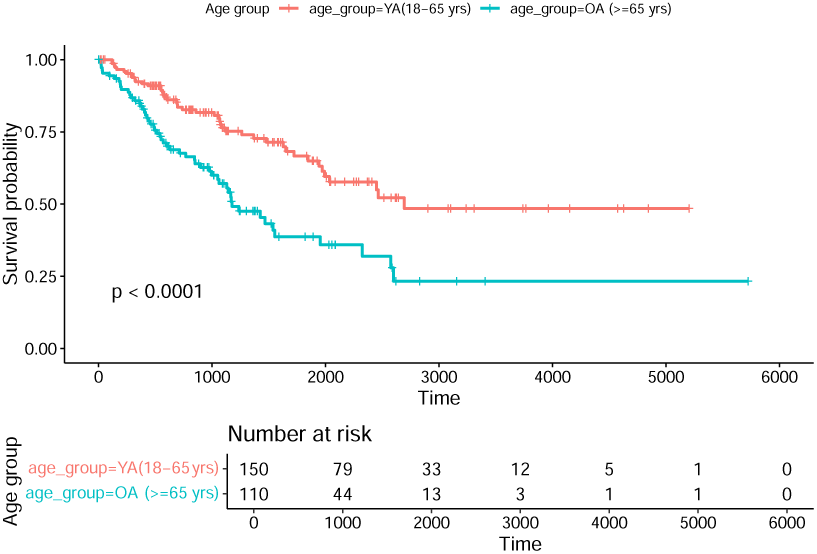
The Kaplan-Meier plot (p-value based on log-rank test) for age-based classification of patients.

## 3 Methodology

### 3.1 Data acquisition and pre-processing

RNA-seq and clinical data from SARC patient samples were acquired from TCGA using the TCGAbiolinks R package [58]. To analyze the gene expression data, we collected unstranded data and performed normalization by gene length and quantile filtered to eliminate low-expression genes using TCGAbiolinks R packages. The samples were grouped as 108 older age (OA) and 154 younger age (YA) patients based on the age of diagnosis ≥ 65 years and 18-65 years respectively. Samples with missing age of diagnosis information were excluded. The overall workflow followed in the analysis is summarized in Fig. 3.1.1.

**Figure 3.1.1.**
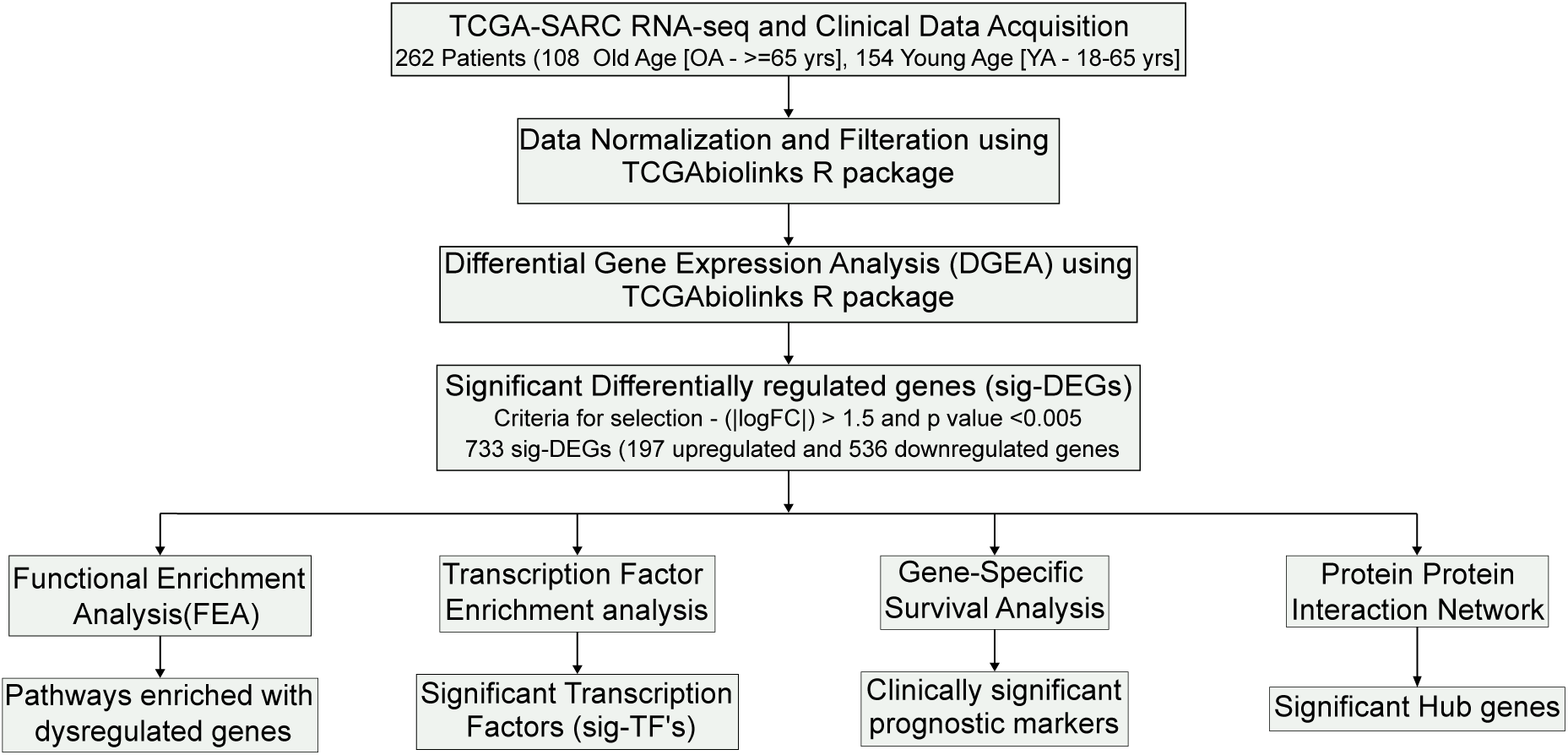
Pipeline followed in the study

### 3.2 Differential gene expression analysis and Functional Enrichment analysis

Differential gene expression analysis (DGEA) was performed using the TCGAanalyze DEA() function (edgeR pipeline) from the TCGAbiolinks R package [58] to compare the gene expression levels between OA and YA age groups. Genes were identified as upregulated, downregulated, or non-significant based on a log fold-change (logFC) threshold of ±1.5 and a false discovery rate (FDR) cut-off *<* 0.005. The Ensembl IDs of the DGEA results were converted to gene symbols using the org.Hs.eg.db R package. An enhanced volcano plot was generated to visualize all the differentially expressed genes (DEGs) using the EnhancedVolcano() function. A heatmap with diverging color palette was generated using the heatmap.2() function to visualize the significant DEGs (sig-DEGs).

Functional enrichment analysis (FEA) and pathway analysis were performed for the upregulated and downregulated genes using the TCGAanalyze EAcomplete() function from the TCGAbiolinks R package [58]. The results of enrichment analysis were presented as bar plots using the TCGAvisualize EAbarplot() function to highlight the top 5 most enriched terms based on fold enrichment and FDR values for biological processes, cellular components, molecular functions and pathways.

### 3.3 Transcription factor enrichment analysis (TFEA)

Significant differentially regulated TFs (sig-TFs) were identified based on transcription factor enrichment analysis and 3 selection criteria, as illustrated in Supplementary Figure 2. Transcription factor databases TRRUST and DoRothEA were used to identify differentially regulated transcription factors (TFs). The dorothea hs database in the dorothea R package [59] contains 1395 TFs 20244 TG’s with 486676 unique interactions, while TRRUST online database [60] contains 9396 interactions of 795 TFs and 2492 TGs. Primarily, comparing the sig-DEGs with online databases, 43 and 21 TFs from the dorothea hs database and TRRUST were seen to be differentially expressed in the OA with respect to YA patient group. For further analysis, the following selection criteria were used. Firstly, TFs that had at least one dysregulated target gene (TG) were selected. Secondly, we negated the interactions where the type of regulation (activation/repression) was not known, and if the type of regulation was known, we selected transcription factors (TFs) with at least one downstream target gene (TG) that exhibited the same expression pattern (up or down-regulation) as the TF in cases of activation, or an opposite expression pattern in cases of repression. Lastly, in the DoRothEA database we selected TFs and TGs having high confidence interaction evidence (A-C). To visualize the interaction between the sig-TF and their TGs, Cytoscape [61] (Available at: Cytoscape) and STRING database [62] (Available at: STRING) was used for network analysis and identifying top 5 associated gene ontology biological process (GO-BP) terms of each sig-TFs. Further, functional enrichment analysis for all the sig-TFs was performed using ShinyGO [63] (Available at: ShinyGO) and represented using R.

### 3.4 Clinical significance of the differentially expressed genes

To assess the impact of the 733 significant DEG’s on survival of the OA group, survival analysis was performed using Cox proportional hazards regression analysis (using survival package in R) on the 108 OA samples stratified by the gene’s expression as followed in methodology by Tanf *et al.,* 2018 [64] and illustrated in Supplementary Figure 3. The expression values of the genes were classified as either high (expression value ≥ median) or low (expression value *<* median). Some of the genes (n=256) were negated from the analysis since the expression values were skewed and their median values were equal to the minimum/maximum/quartile values. Out of the remaining 477 genes, 31 were identified to be significantly associated with patient prognosis (p≤0.05). Further, correlating the gene expression patterns with hazard ratios (HRs) to evaluate the directionality of their impact on survival outcomes, 16 genes were identified to significantly influence the overall survival of the OA group. Further, Kaplan-Meier analysis was used to plot Kaplan-Meier plots for each of the 16 genes along with the p-values for the log-rank association test. Additionally, functional enrichment analysis of the 16 genes was performed using the enrichR package [65] in R.

### 3.5 Network analysis

The protein-protein interaction (PPI) network of 538 out of 733 differentially expressed genes was visualized in Cytoscape version 3.10.2 using the STRING (Search Tool for the Retrieval of Interacting Genes/Proteins) database [66] (Available at: STRING). The network was subjected to ‘Network Analysis’ to obtain the network parameters [67]. Additionally, the top 10 gene hubs were identified using Cytohubba plug-in from the network based on the closeness, degree, and MCC algorithms [68]. The results from these three algorithms were analyzed using a Venn diagram to select the 10 top-ranking genes (Fig. 3.5.1) [69, 70].

**Figure 3.5.1.**
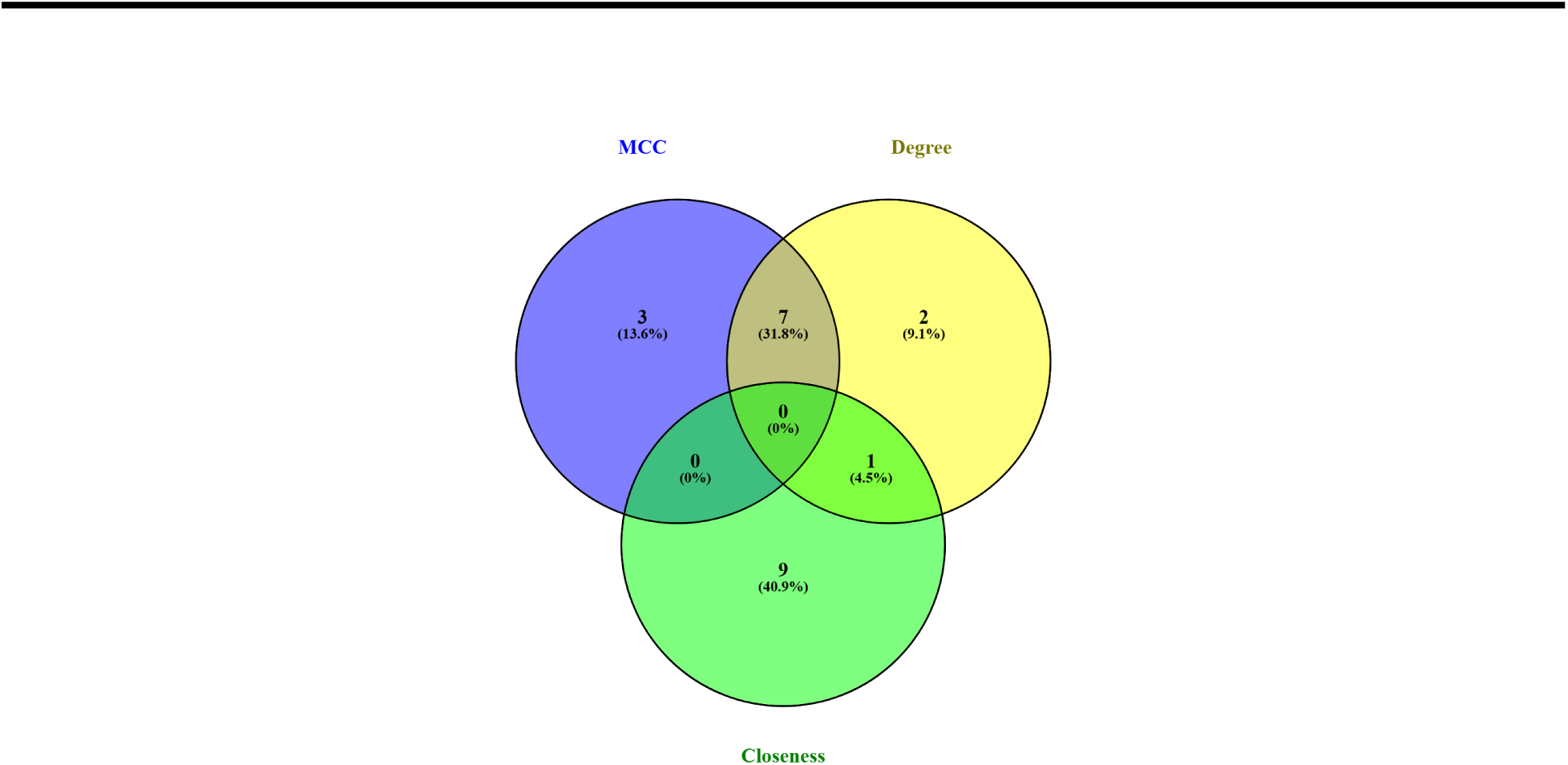
Venn diagram representing intersection of the top 10 ranking genes from the closeness, degree, and MCC algorithms.

## 4 Results and Discussion

### 4.1 Differential gene expression analysis (DGEA)

The DGEA identified 733 significant differentially expressed genes (DEG’s) from the ∼ 23,000 genes, between the two age groups (YA and OA), using a log fold-change and p-value threshold. Of these significant DEG’s, 197 and 536 genes were upregulated and downregulated, respectively in the OA patient group. The volcano plot as depicted in Fig. 4.1.1a shows the distribution of all genes (∼23,000 genes), with significant genes (sig-DEG’s) displayed in the top right and left quadrants. In addition, the top 10 dysregulated (up/down regulated) genes are also highlighted in the plot. The heatmap of the significant DEGs (733 genes), as illustrated in (Fig. 4.1.1b), shows a pattern of dysregulation specific to each age group as clustered by column color (red-OA and blue-YA patient group).

**Figure 4.1.1.**
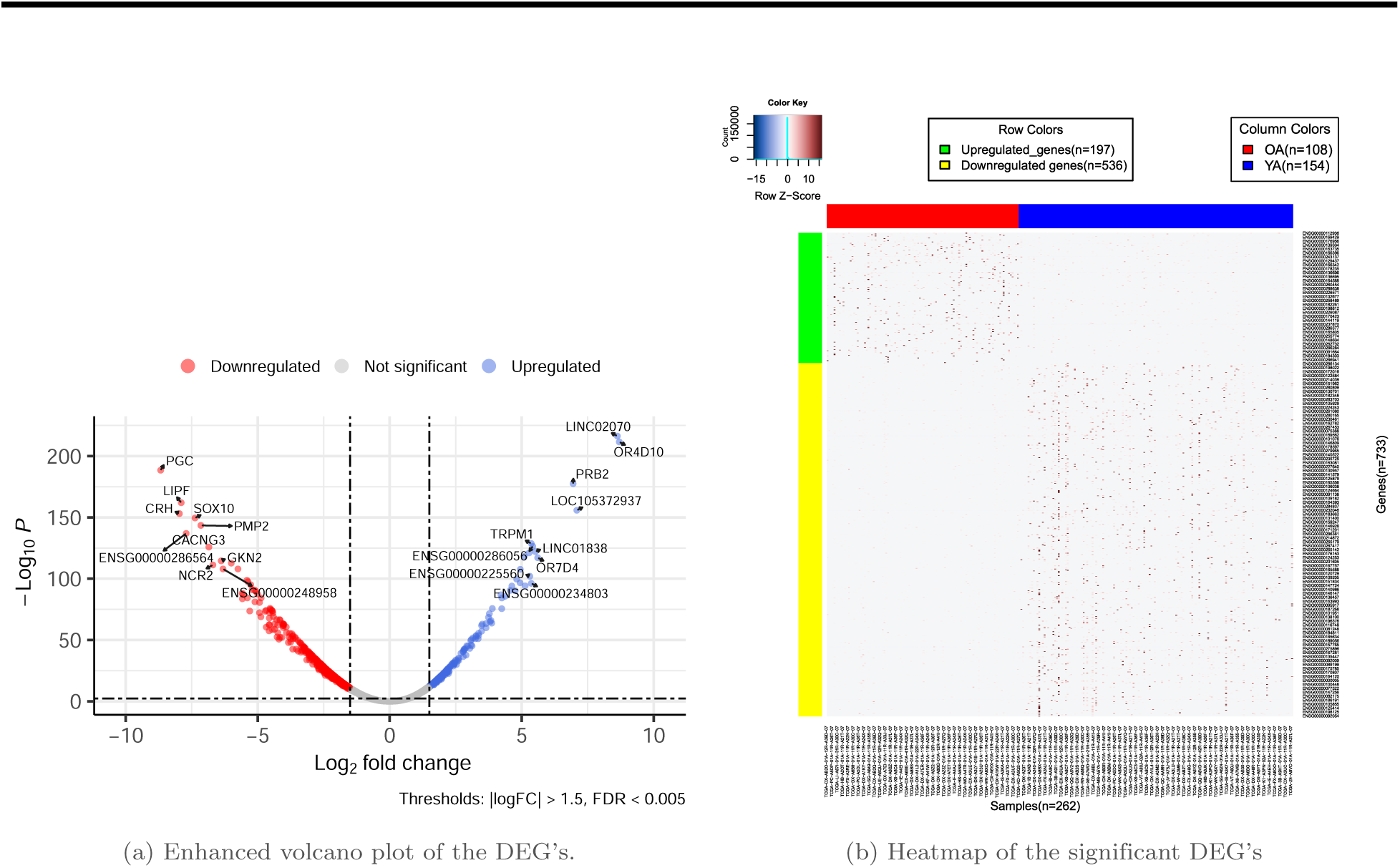
Differential gene expression analysis between OA and YA sarcoma patients (a) Volcano plot (b) Heatmap

The top 10 significantly upregulated genes included 4 protein coding genes (*OR4D10, PRB2, OR7D4, TRPM1*) and 6 lncRNAs (*LINC02070, LOC105372937, LINC01838, ENSG00000286056, ENSG00000234803 (FAM197Y2), ENSG00000225560(FAM197Y8)*). The identified lncRNA’s role in cancer has not yet been characterized. *OR4D10* and *OR7D4*, which are olfactory receptors (OR), are GPCRs mainly involved in the transduction of odorant signals. OR genes have been reported to regulate cell migration, tumor proliferation, secretion, and apoptosis [71], and was found to be upregulated in breast cancer, promoting tumor formation in the same [72]. Further, some OR genes were established as biomarkers for prostate cancer and lung cancer [73]. In the current study, *OR4D10* and *OR7D4* genes which have not yet been explored for their role in cancer, were significantly upregulated in the OA group, suggesting they can be novel potential biomarkers for the same. Higher expression of *TRPM1* (Transient Receptor Potential Cation Channel Subfamily M Member 1) was reported to promote tumor progression in melanoma by activating calcium signaling (Ca2+/calmodulin-dependent protein kinase II*δ* (CaMKII*δ*)/AKT pathway), and was associated with shorter survival in acral melanoma [74]. Notably, the upregulated *PRB2* (Proline Rich Protein BstNI Subfamily 2), which belongs to the retinoblastoma tumor suppressor gene family, codes for a protein involved in inhibiting proliferation through cell cycle control. This gene is established as a TSG in multiple cancers [75, 76] and its downregulation is also associated with lower survival in sarcoma [77].

The top 10 significantly downregulated genes included 8 protein coding genes (*PGC, CRH, LIPF, SOX10, PMP2, CACNG3, NCR2, GKN2*) and 2 uncharacterized lncRNA’s (*ENSG00000286564, ENSG00000248958*).

*PGC* (Progastricsin) is a gene involved in protein digestion and has shown abnormal expression in cancers such as gastric and breast cancers [78, 79]. It acts as a tumor suppressor by inhibiting proliferation and invasion of gastric cancer cells, promoting favorable immune infiltration, ECM modification and is associated with a good patient prognosis in multiple other cancers [80–82]. Notably, the downregulated *CRH* (Corticotropin releasing hormone) gene was shown to regulate stress and immune response [83, 84], and acts as an oncogene by promoting tumor immunoescape and angiogenesis in multiple cancers such as breast, ovarian, and colon cancer [85–87]. *LIPF* (Lipase F, gastric type) gene, involved in digestion processes, is reported to be downregulated in gastric cancer and is associated with a poor prognosis in the same [88, 89]. Since liposarcoma (a subtype of STS) is associated with altered lipid metabolism [90], the downregulation of *LIPF* can potentially influence tumor growth in the same way. *SOX10* (SRY-Box transcription factor 10) gene exhibits a dual role in the context of cancer [91], acting as a tumor suppressor via the Wnt/*β*-catenin pathway inhibition [92–94], and an oncogene via promoting PI3K/AKT pathways [95]. *SOX10* is also commonly associated with neuroblastoma, melanoma, and is a diagnostic marker in sarcoma [96–98]. *PMP2* (Peripheral Myelin Protein 2), a gene that is activated by *SOX10* and in turn promotes melanoma cell invasion in some subsets [99], was also downregulated in OA patients similar to its upstream controller gene. Low expression of the downregulated *CACNG3* (Calcium Voltage-Gated Channel Auxiliary Subunit Gamma 3), a gene associated with brain tumors, was reported to be linked to shorter survival, suggesting a tumor suppressive role [100]. Further, the downregulated *NCR2* (Natural Cytotoxicity Triggering Receptor 2) is a gene involved in activation of the innate immune response and tumor recognition [101, 102]. Another significant gene identified is the downregulated *GKN2* (Gastrokine-2), which is reported to be associated with preventing inflammation, proliferation, migration, and metastases in gastric and lung cancers [103–105]. Additionally, it is a potential clinical biomarker associated with ECM degradation and a lower prognosis in gastric and lung cancer [104, 105].

These findings show that except for *CRH* and *PMP2*, the identified downregulated genes *PGC, LIPF, SOX10, CACNG3, NCR2* and *GKN2* could have a tumor suppressive role in OA sarcoma patients similar to other cancers, impacting multiple pathways involved in immune response, ECM modifications, lipid metabolism, Wnt/*β*-catenin and resulting in their lower prognosis. The upregulated *TRPM1* gene was reported to play an oncogenic role in multiple cancers by promoting calcium signaling (Ca2+/calmodulin-dependent protein kinase II*δ* (CaMKII*δ*)/AKT pathway), and could have a similar role in OA sarcoma patients. Additionally, many dysregulated lncRNA’s and genes identified in the study, are yet to be characterized for their biological significance in cancer, paving a way for new research.

Functional enrichment analysis (FEA) revealed an enrichment of dysregulated genes in MMP (Matrix metalloproteinases) inhibition pathways, calcium signaling, tight junction signaling and Integrin-Linked Kinase (ILK) signaling. These signaling pathways are associated with ECM remodeling and mediating the metastases process in cancer cells. MMPs promote tumor proliferation by degrading matrix barriers such as the ECM and enhancing angiogenesis [106, 107]. Calcium signaling is associated with EMT, promoting cell migration and metastasis [108]. Tight junction proteins are downregulated during EMT causing decreased cell-cell adhesion, thus enhancing cell motility and tumor invasiveness [109]. ILK (Integrin-linked kinase) signaling is involved in the regulation of cell proliferation, migration, differentiation, and survival [110]. These observations highlight that the significant differentially expressed genes in the OA group alter the tumor microenvironment and control ECM interactions. The top 5 GO terms and KEGG pathways enriched with the 197 upregulated and 536 downregulated are represented in Fig. 4.1.2a and Fig. 4.1.2b respectively.

**Figure 4.1.2.**
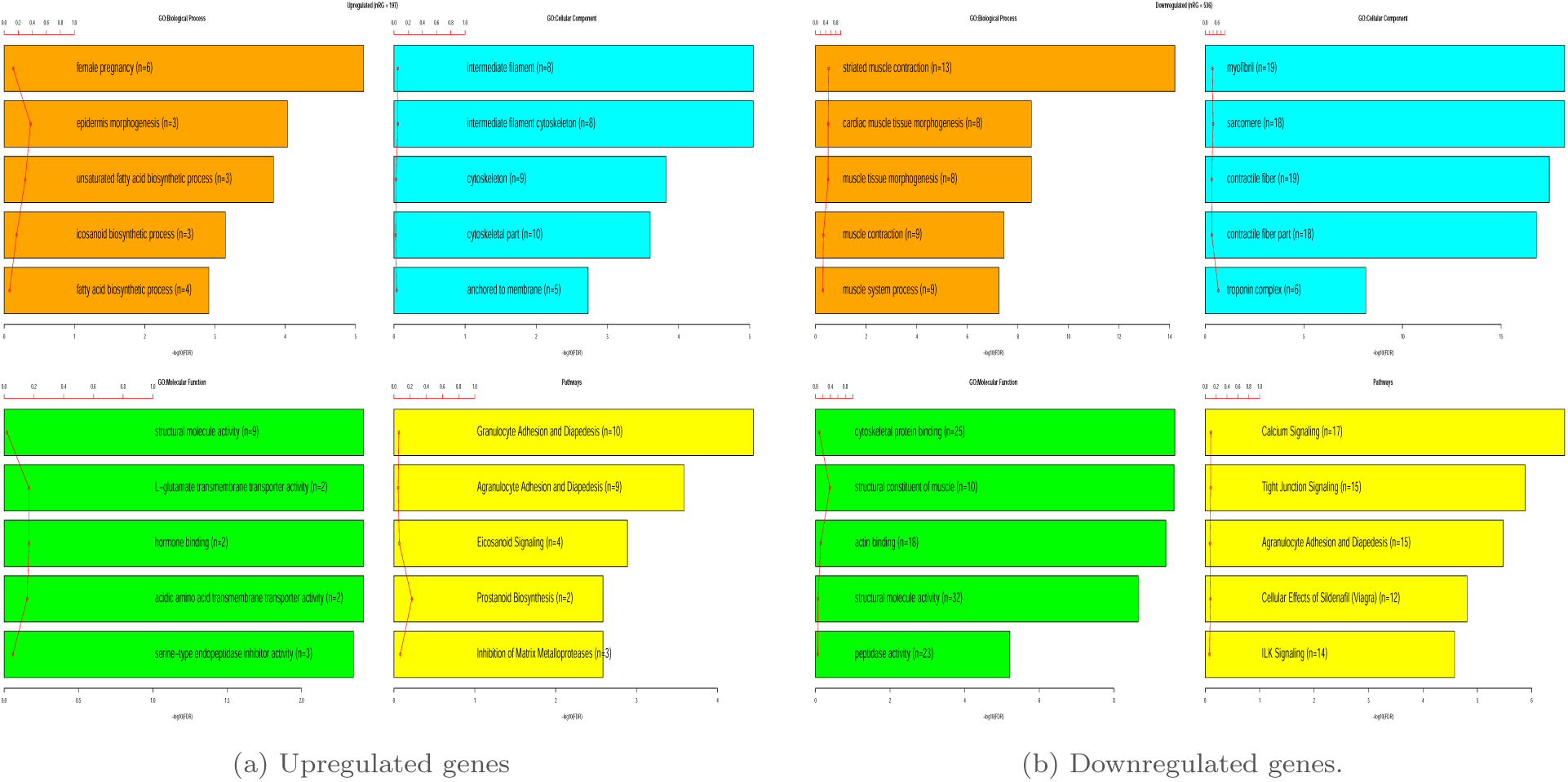
Functional enrichment analysis and pathway analysis.

### 4.2 Transcription factor enrichment analysis (TFEA)

Transcription factors are master biological molecules that control the gene expression of multiple genes and hence regulate multiple downstream biological processes and undergo differential expression in cancer [111]. In this study, we narrowed down 43 and 21 TFs that are differentially regulated in the OA compared to the YA group from the TF databases DoRothEA and TRRUST, respectively. Based on three selection criteria as explained in the methodology, 10 sig-TFs were identified, namely *CDX2, ELF3, GRHL2, HNF4A, OXT2, SOX10, NR5A1, SPDEF, PHOX2B,* and *PGR*. The TF’s *CDX2, HNF4A,* and *SOX10* were present in both the transcription factor databases-based analyses. Though *ELF3* was detected in both databases-based analyses, it was negated in the TRRUST database-based analysis since the interaction with its target gene was unknown in the same. Moreover, it is to be noted that more transcription factors and interacting TGs were identified from the DoRothEA database compared to TRRUST, this could be due to the smaller size of the database.

Using the STRING interaction database and Cytoscape tool the interactions between the differentially regulated TGs and sig-TF along with the biological process (BP) GO term they are associated with is illustrated in Fig. 4.2.1 (c) and (d). Fig. 4.2.1 (b) represents the top 5 GO terms enriched with the 10 sig-TFs for cellular component (CC), molecular function(MF), and biological process (BP). The most enriched biological process term is epithelial cell differentiation (log_2_(fold enrichment)) and most of them (9 out of 10) are associated with “Positive regulation of transcription by RNA polymerase II” since they are transcription factors.

**Figure 4.2.1.**
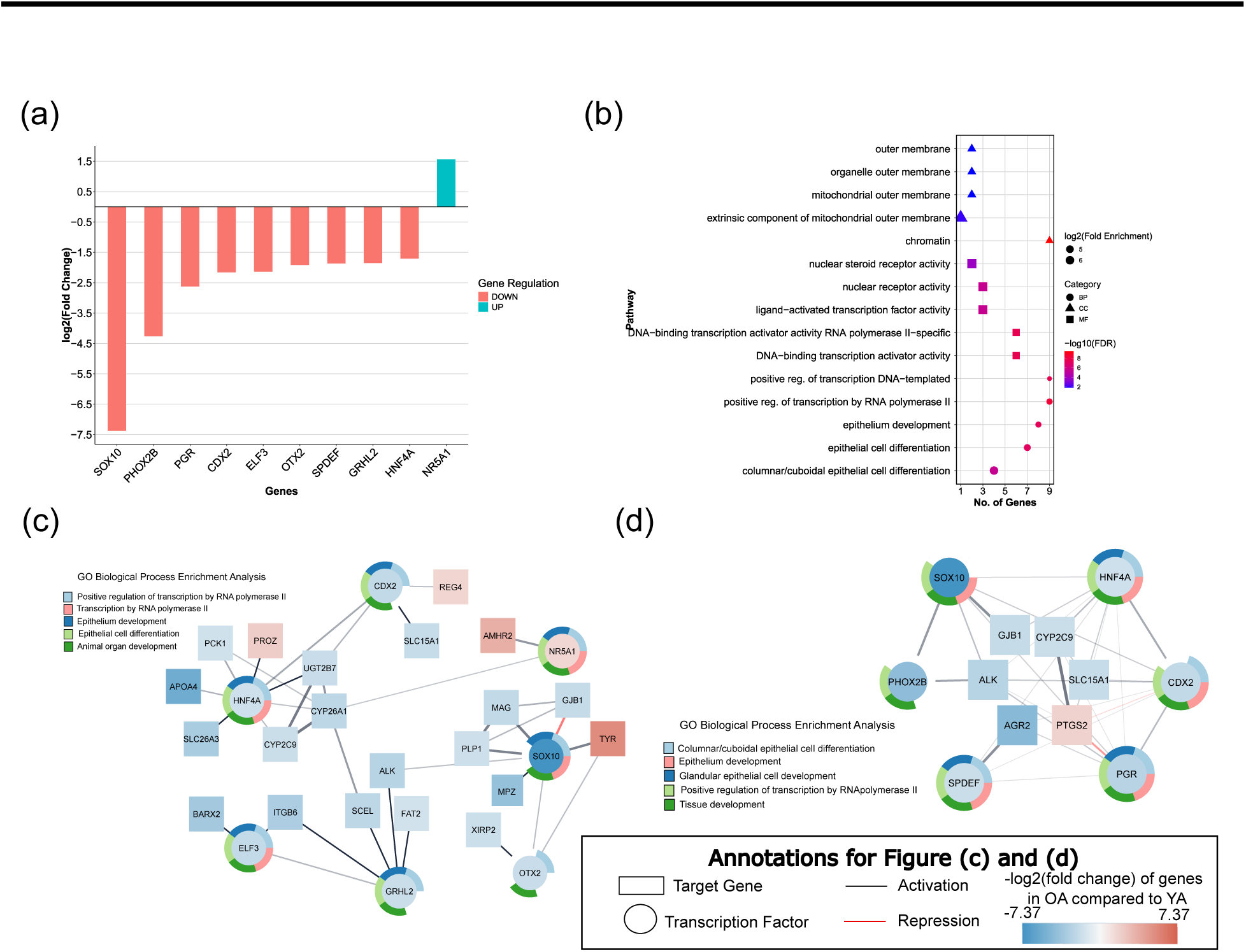
Gene expression analysis, Functional Enrichment analysis and interactions of significant differentially expressed transcription factors (sig-TF; n=10). (a) Bar graph of the log fold-change in expression of the sig-TF between the OA and YA group. (b) Functional enrichment analysis of the 10 sig-TF. (c) Interaction of the 7 sig-TF and their differentially regulated target genes (TG; n=24) using DoRothEA database and STRING Network. (d) Interaction of the sig-TF (n=6) and their differentially regulated target genes (TG; n=6) using TRRUST database and STRING Network.

Among the sig-TFs, only *NR5A1* was upregulated in the OA group while the others were downregulated as shown in Fig. 4.2.1 (a). The only upregulated gene *NR5A1* is not well studied in cancer, however it is involved in promoting autonomous self-renewal of pluripotent stem cells, indicating a possibility of an oncogenic role and requires further investigation [112]. Although some of the downregulated sig-TFs have dual roles in different cancers, they have tumor suppressive roles and their downregulation is associated with a lower prognosis paving ways to explore their role in the progression of sarcoma in older age group.

Table 4.2.1 provides a brief description of current literature studies on the sig-TF and their biological significance. Among these, the TFs *GRHL2, ELF3, PGR* and *CDX2* which are downregulated in our study, were previously reported to have tumor suppressive roles in multiple types of sarcoma by promoting epithelial characteristics and inhibiting EMT, indicating a similar role in OA sarcoma patients [113–125]. However, the *SOX10* transcription factor has a tissue-specific role in different cancers. This gene was previously reported to have an oncogenic role (promoting stem-like properties) and a diagnostic marker for malignant peripheral nerve sheath tumor (MPNST) and Synovial sarcoma, was downregulated in the OA patients [126]. This suggests the possibility of tumor suppressive role of *SOX10* in OA patients, which needs further validation. Though the role of downregulated *HNF4A, SPDEF, PHOX2B* and *OXT2* have not been explored in sarcoma, except *OXT2* having an oncogenic role [127, 128]), others have shown a tumor suppressive role in other cancers as summarized in Table 4.2.1 [129–131] Hence, proposing a TSG role for the TFs *HNF4A, SPDEF*, and *PHOX2B* in OA sarcoma patients as well.

**Table 4.2.1.** Role of sig-TF in cancer and sarcomas.

| Transcription Factor | Regulation in OA group | Role of gene | References (PMID) |
| --- | --- | --- | --- |
| *GRHL2* (Grainyhead Like Transcription Factor 2) | DOWN | Dual role in cancer, Promotes epithelial like state in sarcoma, associated with increased overall survival in STS<br>Oncogene by facilitating tumorigenesis and metastasis | 27402864 [119],<br>23814079 [132],<br>32194886 [113],<br>35269877[114] |
| *NR5A1* (Nuclear Receptor Subfamily 5 Group A Member 1) | UP | Promote MAPK independent autonomous cell self-renewal | 32084386 [112] |
| *HNF4A* (Hepatocyte Nuclear Factor 4 Alpha) | DOWN | Dual role in cancer- TSG role by suppressing EMT via inhibition of β-catenin.<br>Oncogenic role by promoting angiogenesis | 31695151 [129],<br>25592038 [133] |
| *ELF3* (ETS-Related Transcription Factor Elf-3) | DOWN | Dual role in cancer - TSG role as regulator of epithelial differentiation inhibits EMT progression, associated with Synovial Sarcoma.<br>In NSCLC and neuroendocrine tumors, it activates downstream oncogenic targets. | 12414507 [115],<br>32483824 [116],<br>17060315 [117],<br>36864480 [134],<br>31780666 [135],<br>36840413[136] |
| *SOX10* (SRY-Box Transcription Factor 10) | DOWN | Diagnostic biomarker for subtypes of sarcoma (malignant peripheral nerve sheath tumor)<br>TSG role in gastrointestinal cancers via inhibiting EMT and Wnt/β-catenin signaling<br>Oncogenic role by promoting migration in melanoma | 25301735 [92],<br>32669612 [94],<br>23929265 [98],<br>32592167 [137],<br>25724000 [96],<br>38132479 [138] |
| *CDX2* (Caudal Type Homeobox 2) | DOWN | Promotes epithelial differentiation (loss of epithelial differentiation observed in *CDX2* absent MPNST)<br>TSG role via inhibition of proliferation by suppressing Wnt/β-catenin signaling | 35122289 [120],<br>35328309 [121],<br>32117703 [122],<br>33239678 [123],<br>30631044 [139],<br>34616012 [125] |
| *OTX2* (Orthodenticle Homeobox 2 ) | DOWN | *OTX2* is a developmentally regulated transcription factor.<br>Oncogenic role in medulloblastoma by promoting metastases via mTOR signaling | 38674001 [127];<br>20028867 [128] |
| *SPDEF* (SAM Pointed Domain Containing ETS Transcription Factor) | DOWN | Dual role in cancer.<br>Involved in the regulation of growth, proliferation, migration in different contexts | 32547225 [130],<br>35951453 [140],<br>34667150 [141] |

| Transcription Factor | Regulation in OA group | Role of gene | References(PMID) |
| --- | --- | --- | --- |
| *PHOX2B* (Paired Like Homeobox 2B) | DOWN | Diagnostic immunomarker for neuroblastoma.<br>Downregulation via promoter methylation promotes metastases in neuroblastoma | 26840262 [131],<br>28640941 [142] |
| *PGR* (Progesterone Receptor) | DOWN | Hormone receptor with a dual role in cancer.<br>Activates tumor progression via macropinocytosis in pancreatic ductal cancer<br>Acts as a tumor suppressor in cervical cancer In STS male patients, *PGR* negative expression was associated with low prognosis | 33139507[143],<br>25229191 [144],<br>23497154 [118] |

Function enrichment analysis revealed that 8 of the sig-TF’s are involved in epithelial development (*GRHL2, ELF3, SOX10, HNF4A, NR5A1* and *CDX2*). Supporting this, previous studies have also reported the role of *GRHL2, ELF3, HNF4A, SOX10,* and *CDX2* in promoting epithelial differentiation and inhibiting EMT (epithelial-mesenchymal transition) progression and Wnt/*β*-catenin signaling pathway in cancers and in specific sarcomas as well [92, 113, 115–117, 119–123, 125, 129, 134, 139]). Further, increased expression of *GRHL2* was associated with increased metastases free and improved overall survival in sarcoma [119]. These findings indicate a possible loss of epithelial state in OA patients, which in turn could lead to the initiation of EMT and activation of the Wnt/*β*-catenin signaling pathway, which needs to be further validated.

### 4.3 Clinical significance of the differentially expressed genes

To obtain a list of genes that have a prognostic significance on the OA group, survival analysis was performed using univariate Cox proportional hazards regression analysis on the 477 sig-DEGs. The analysis revealed the presence of 31 genes that had a significant association with prognosis (p≤0.05). However, correlating each gene’s expression group (High/ Low expression group) that was associated with the *>*1 hazard ratio (poorer prognosis) and their mRNA regulation (UP/DOWN) status in the OA group, 16 genes (referred to as sig-Survival genes) were identified that were used for further analysis. Fig. 4.3.1 (b) represents the log fold-change expression of the sig-Survival genes in the OA compared to the YA group. Similar to the sig-TF’s, most of the identified genes (n=13) were downregulated in the OA group. Fig. 4.3.1 (a) and Fig. 4.3.2 represent the K-M plot of the 16 genes obtained by KM analysis and HR, 95%-CI, and p-values of sig-Survival genes in the OA group obtained from Cox regression analysis, respectively.

**Figure 4.3.1.**
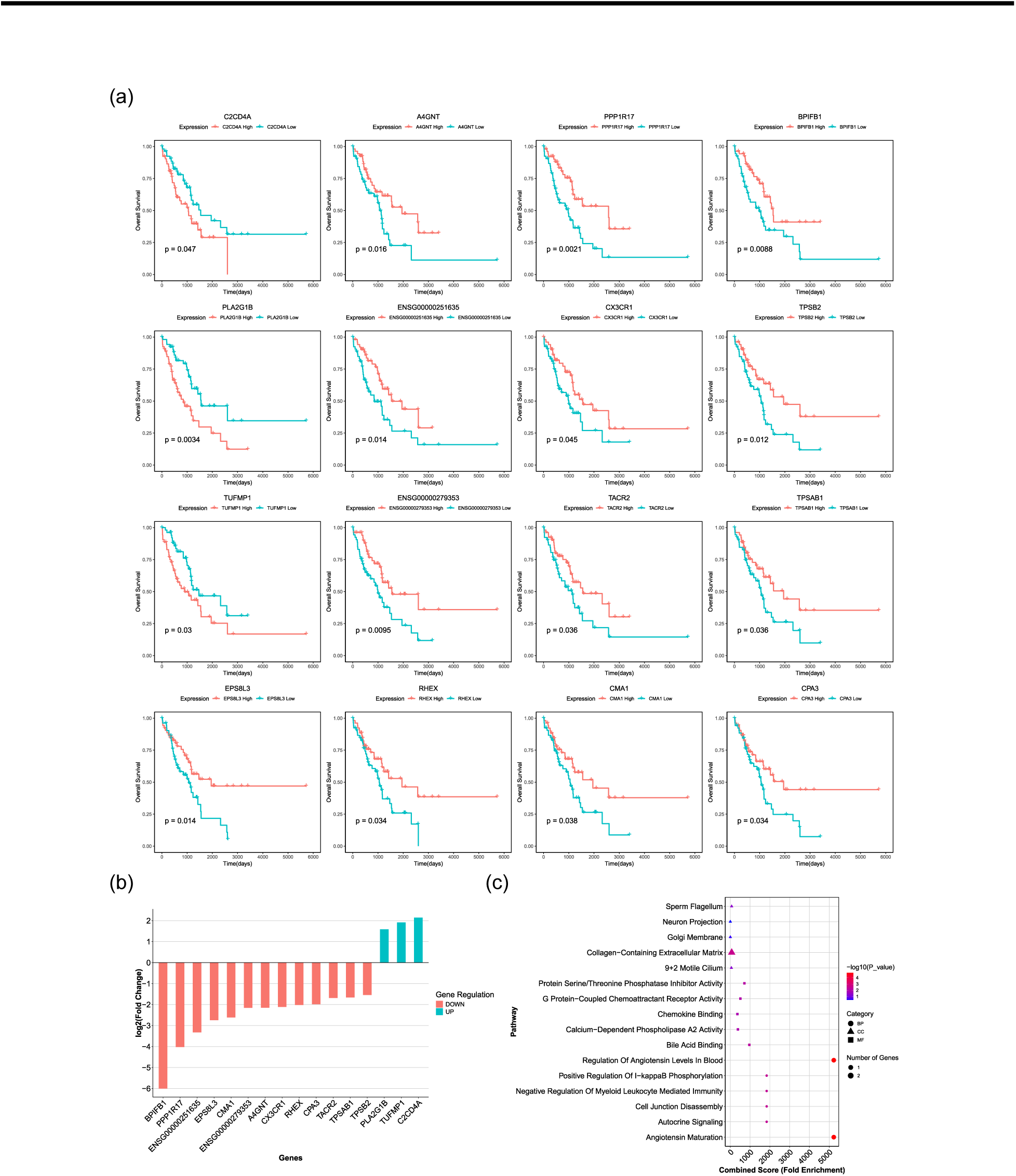
Univariate survival based on DEG’s expression stratification, their log fold-change expression and functional enrichment analysis. (a) The Kaplan-Meier plot (p-value based on log-rank test) of 16 genes (sig-Survival) that showed a significant association with the prognosis of patients when stratified by their expression. (b) The log fold-change in mRNA expression of the 16 genes in the OA group with respect to the YA group (DGEA results). (c) Bubble plot representing the functional enrichment analysis of the 16 genes representing the top 5 gene ontology terms for cellular component, molecular function and biological process.

**Figure 4.3.2.**
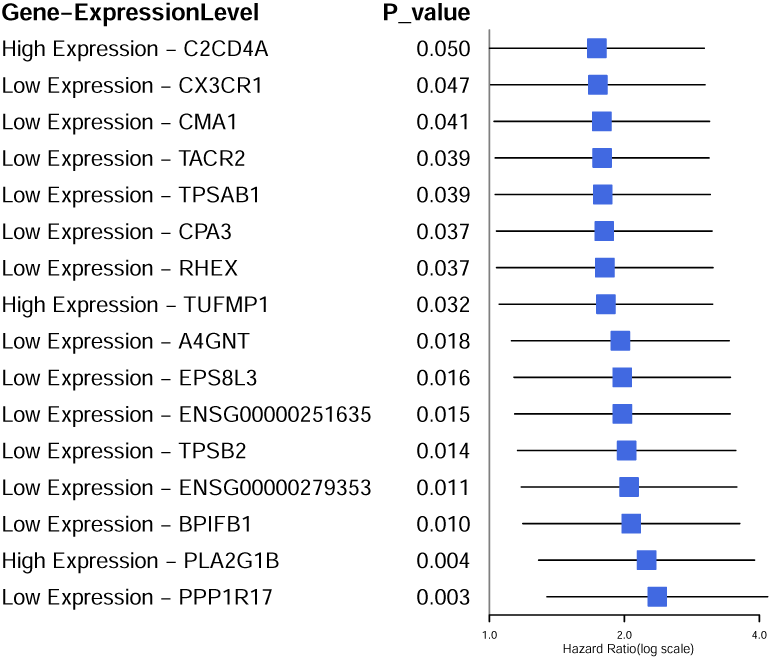
Univariate Cox regression analysis results of the 16 significant genes identified.

None of the 16 identified genes had been previously reported in sarcoma, furthermore, there were no reported studies for 5 of these genes in cancer (*TUFMP1, TPSAB1, ENSG00000279353, TPSB2* and *ENSG00000251635*). Some of these genes have been associated with prognosis in several other cancers, as summarized in Table 4.3.1 along with the role of these genes in cancer.

**Table 4.3.1.**
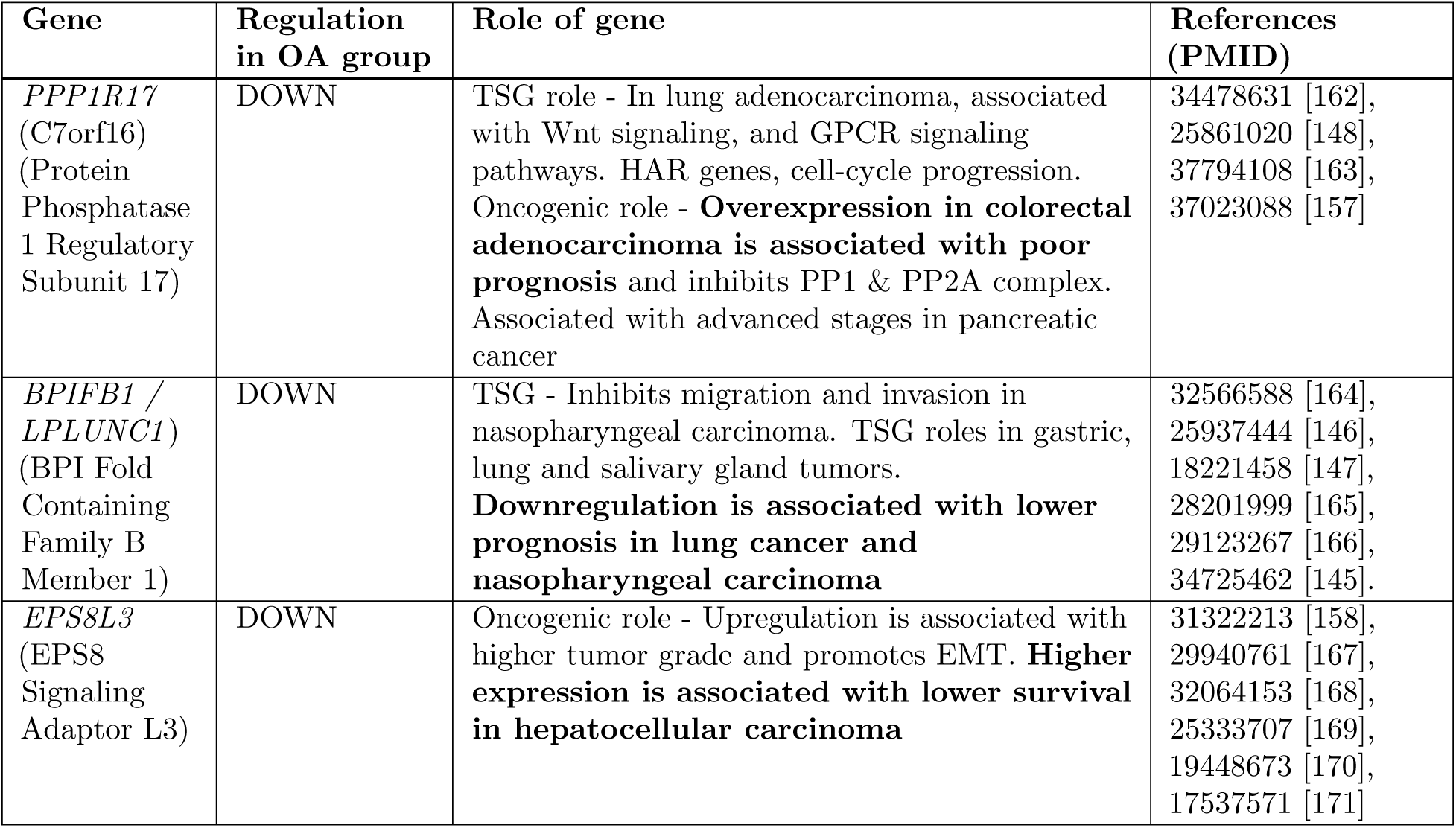
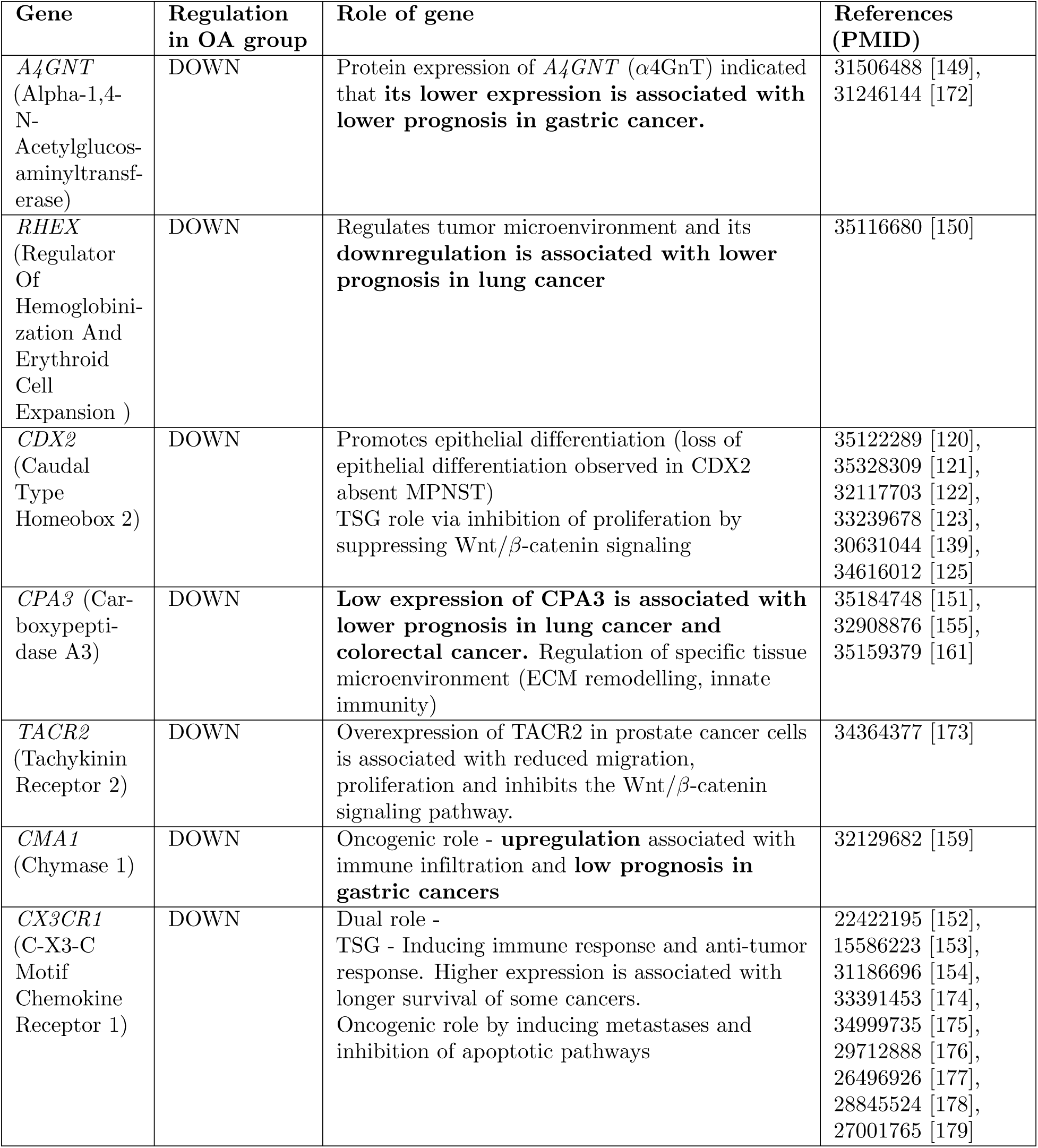
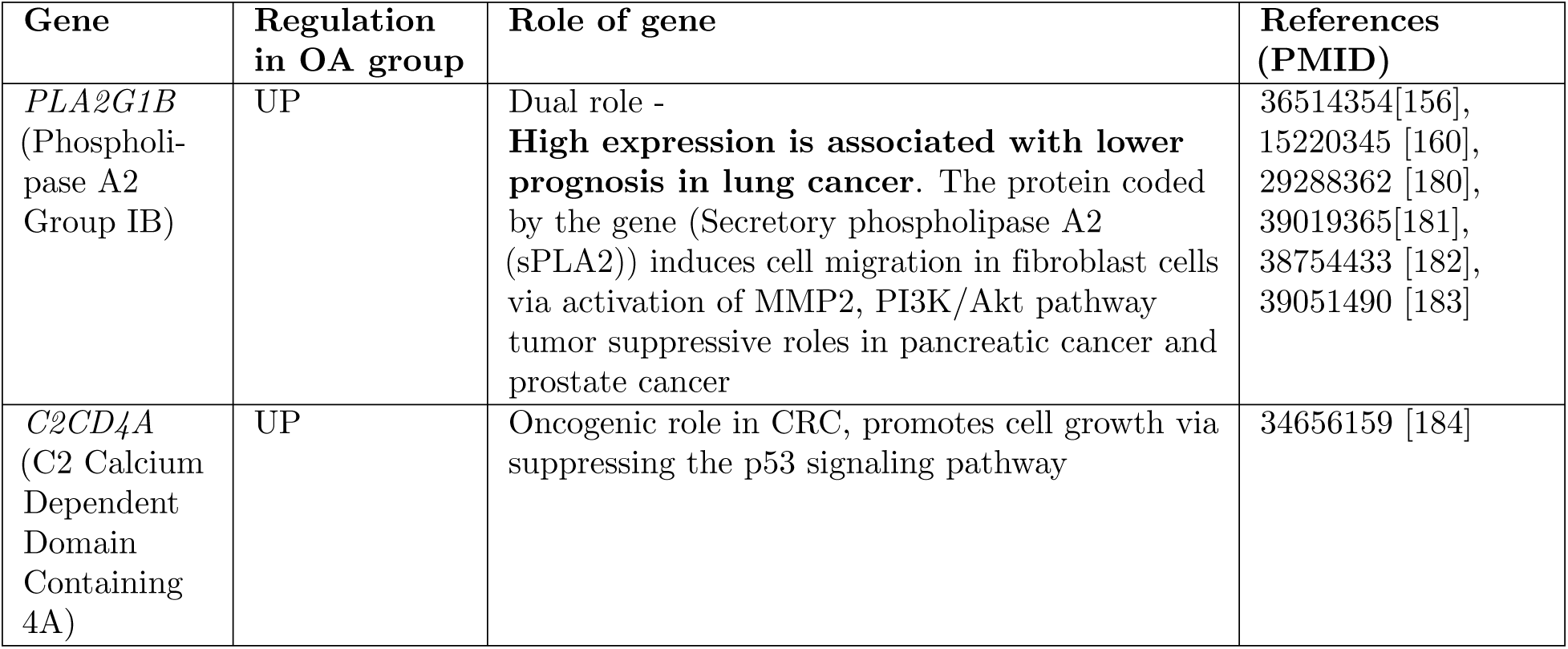
Role of identified potential prognostic biomarkers (OA group) in cancer.

| Gene | Regulation in OA group | Role of gene | References (PMID) |
| --- | --- | --- | --- |
| *PPP1R17* (C7orf16) (Protein Phosphatase 1 Regulatory Subunit 17) | DOWN | TSG role - In lung adenocarcinoma, associated with Wnt signaling, and GPCR signaling pathways. HAR genes, cell-cycle progression. Oncogenic role - **Overexpression in colorectal adenocarcinoma is associated with poor prognosis** and inhibits PP1 & PP2A complex. Associated with advanced stages in pancreatic cancer | 34478631 [162], 25861020 [148], 37794108 [163], 37023088 [157] |
| *BPIFB1* / *LPLUNC1* (BPI Fold Containing Family B Member 1) | DOWN | TSG - Inhibits migration and invasion in nasopharyngeal carcinoma. TSG roles in gastric, lung and salivary gland tumors. **Downregulation is associated with lower prognosis in lung cancer and nasopharyngeal carcinoma** | 32566588 [164], 25937444 [146], 18221458 [147], 28201999 [165], 29123267 [166], 34725462 [145]. |
| *EPS8L3* (EPS8 Signaling Adaptor L3) | DOWN | Oncogenic role - Upregulation is associated with higher tumor grade and promotes EMT. **Higher expression is associated with lower survival in hepatocellular carcinoma** | 31322213 [158], 29940761 [167], 32064153 [168], 25333707 [169], 19448673 [170], 17537571 [171] |
| *A4GNT*<br>(Alpha-1,4-N-Acetylglucosaminyltransferase) | DOWN | Protein expression of *A4GNT* (α4GnT) indicated that **its lower expression is associated with lower prognosis in gastric cancer.** | 31506488 [149],<br>31246144 [172] |
| *RHEX*<br>(Regulator Of Hemoglobini-zation And Erythroid Cell Expansion ) | DOWN | Regulates tumor microenvironment and its **downregulation is associated with lower prognosis in lung cancer** | 35116680 [150] |
| *CDX2*<br>(Caudal Type Homeobox 2) | DOWN | Promotes epithelial differentiation (loss of epithelial differentiation observed in CDX2 absent MPNST)<br>TSG role via inhibition of proliferation by suppressing Wnt/β-catenin signaling | 35122289 [120],<br>35328309 [121],<br>32117703 [122],<br>33239678 [123],<br>30631044 [139],<br>34616012 [125] |
| *CPA3* (Carboxypeptidase A3) | DOWN | **Low expression of CPA3 is associated with lower prognosis in lung cancer and colorectal cancer.** Regulation of specific tissue microenvironment (ECM remodelling, innate immunity) | 35184748 [151],<br>32908876 [155],<br>35159379 [161] |
| *TACR2*<br>(Tachykinin Receptor 2) | DOWN | Overexpression of TACR2 in prostate cancer cells is associated with reduced migration, proliferation and inhibits the Wnt/β-catenin signaling pathway. | 34364377 [173] |
| *CMA1*<br>(Chymase 1) | DOWN | Oncogenic role - **upregulation** associated with immune infiltration and **low prognosis in gastric cancers** | 32129682 [159] |
| *CX3CR1*<br>(C-X3-C Motif Chemokine Receptor 1) | DOWN | Dual role -<br>TSG - Inducing immune response and anti-tumor response. Higher expression is associated with longer survival of some cancers.<br>Oncogenic role by inducing metastases and inhibition of apoptotic pathways | 22422195 [152],<br>15586223 [153],<br>31186696 [154],<br>33391453 [174],<br>34999735 [175],<br>29712888 [176],<br>26496926 [177],<br>28845524 [178],<br>27001765 [179] |
| *PLA2G1B*<br>(Phospholipase A2 Group IB) | UP | Dual role - **High expression is associated with lower prognosis in lung cancer.** The protein coded by the gene (Secretory phospholipase A2 (sPLA2)) induces cell migration in fibroblast cells via activation of MMP2, PI3K/Akt pathway tumor suppressive roles in pancreatic cancer and prostate cancer | 36514354 [156], 15220345 [160], 29288362 [180], 39019365 [181], 38754433 [182], 39051490 [183] |
| *C2CD4A*<br>(C2 Calcium Dependent Domain Containing 4A) | UP | Oncogenic role in CRC, promotes cell growth via suppressing the p53 signaling pathway | 34656159 [184] |

Supporting the observations made in this study, downregulation of *BPIFB1, A4GNT, RHEX, CPA3,* and *CX3CR1* genes are shown to be associated with lower prognosis in some cancers [145–155], and upregulation of *PLA2G1B* was shown to be associated with lower prognosis in lung cancer [156]). This indicates that the downregulated genes could act as TSG, while the upregulated genes act as oncogenes in sarcoma, affecting the patient’s prognosis. However, in contradiction to the current findings, higher expression of the downregulated *PPP1R17, EPS8L3*, and *CMA1* genes were previously reported to be associated with lower prognosis in different cancers [157–159]. Nevertheless, many of these genes have both a tumor suppressive (TSG) or an oncogenic role depending on the cancer site, as described in Table 4.3.1. Hence, it is important to conduct studies that explore the role of these genes in sarcoma.

Functional enrichment analysis revealed that the enriched GO terms are involved in angiotensin maturation, extracellular matrix interactions, and immune-mediated pathways (Fig. 4.3.2 (c)). Previous studies also show that the genes *RHEX, CX3CR1, CMA1* are associated with immune regulation pathways [150–152, 159].

*PLA2G1B* was previously reported to impact the extracellular matrix [160]. Additionally, *CPA3* is established to have a role in angiotensin maturation, a process involved in angiogenesis and ECM remodeling which is highlighted in the functional enrichment analysis [161]. These suggest a possible difference in the tumor microenvironment among the older age group that could be responsible for their lower prognosis.

Furthermore, similar to the roles of some sig-TF’s and top 10 up/down-regulated genes which were involved in the Wnt/*β*-catenin signaling pathway and EMT pathways, some of the prognostic genes such as *PPP1R17, BPIFB1, EPS8L3,* and *TACR2* have been reported to impact these pathways. Though the downregulation of the tumor suppressors *PPP1R17, BPIFB1,* and *TACR2,* as observed in the study, is associated with increased migration and activation of *β*-catenin signaling thereby promoting tumor progression, surprisingly the upregulation of *EPS8L3* (observed to be downregulated in the OA group) was shown to be associated with promoting EMT in previous studies [148, 166, 169, 173]. These findings suggest an activation of the Wnt/*β*-catenin signaling pathway and EMT process in the OA group that needs further validation.

### 4.4 Network Analysis: Protein-protein interaction (PPI) network construction, hub gene identification and module analysis

Based on the analysis from the STRING database, the PPI network of 538 out of 733 differentially expressed genes was visualized in Cytoscape. The PPI network consisted of 538 nodes and 2070 edges with a 9.370 average number of neighbors per node. Further, the differential expression data was imported for all sig-DEGs into the STRING database and visualized to investigate the sig-DEGs. In the resulting network, green nodes represented upregulated DEGs, and red nodes represented downregulated DEGs as illustrated in Fig. 4.4.1. The top 10 hub genes were identified using Cytohubba plug-in from the network based on closeness and degree which resulted in distinct hub genes. Thus, to validate the findings, we employed another algorithm, MCC. The hub genes are selected based on their high scores in the degree ranking in CytoHubba (Fig. 3.5.1) due to overlaps with both MCC and closeness, namely *MYH2, MYH7, MYL1, MYL2, MYL11, ACTN2, TTN, TNNI2, MB* and *CXCL8* as shown in Fig. 4.4.2. The data regarding the rank of each gene in each algorithm is provided in the Supplementary Table 1.

**Figure 4.4.1.**
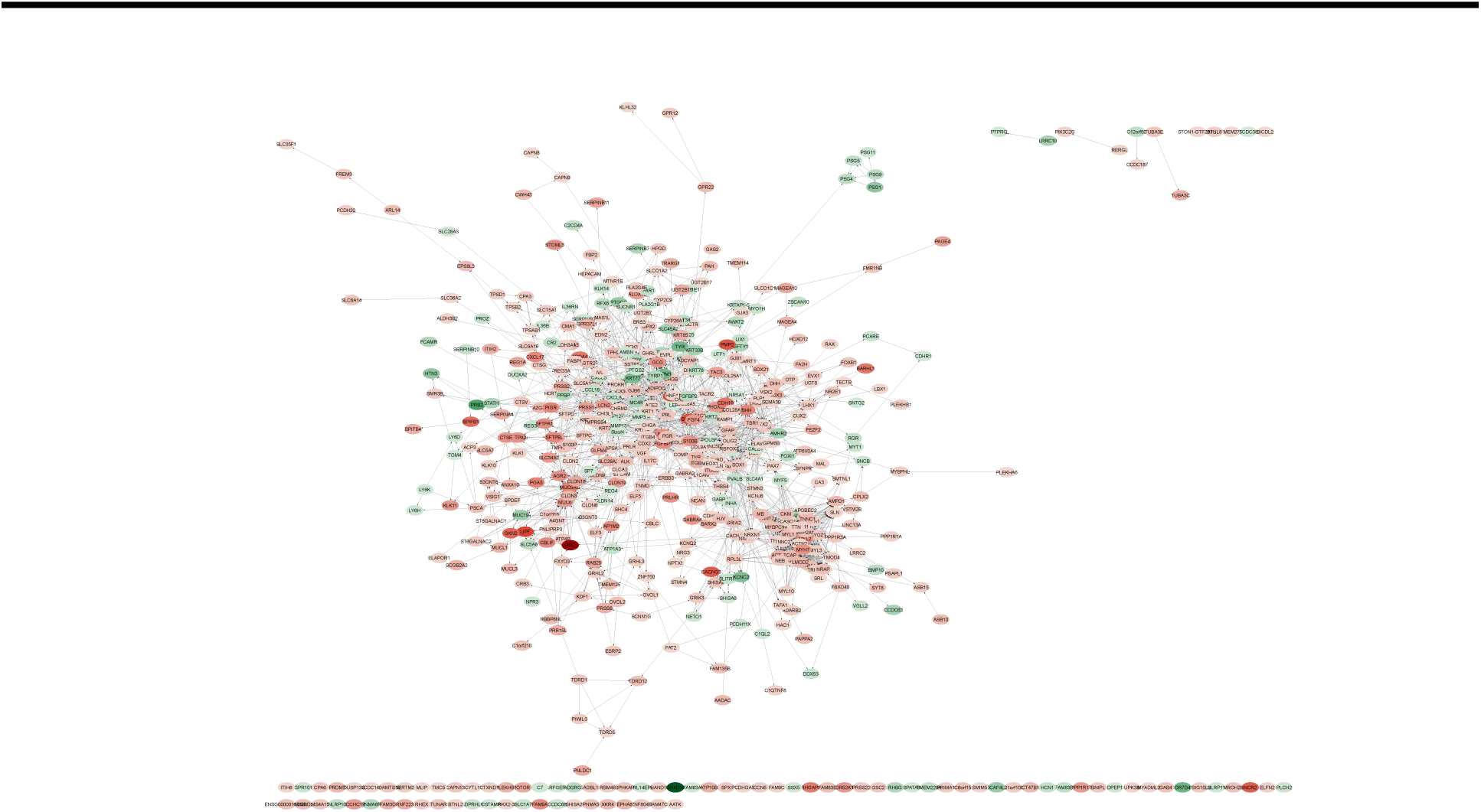
Protein-protein interaction (PPI) network for 538 significant DEGs visualized in Cytoscape software based on the STRING database. Nodes represent DEGs, with green indicating upregulation and red indicating downregulation in older age (OA) patients compared to younger age (YA). Edges represent interaction scores derived from STRING.

**Figure 4.4.2.**
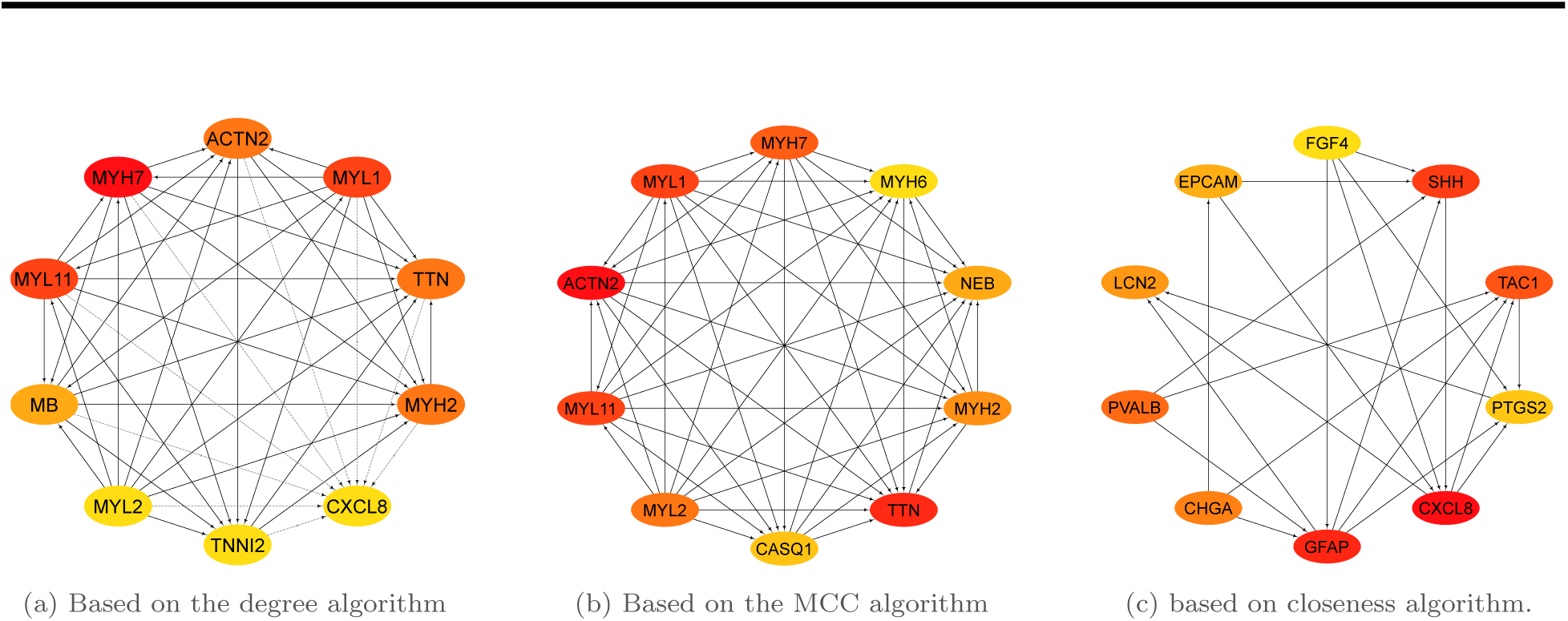
Top 10 hubs identified using Cytohubba in Cytoscape. Nodes represent genes, with red being the highest rank and yellow the lowest, solid lines represent direct link while dotted lines represent indirect links

Myosins, actin-based motor molecules with ATPase activity essential for muscle contraction, constituted 50% of the top 10 hub genes and are all downregulated in this study. Studies showed that the higher expression of some of these myosin genes *MYL2, MYL1, MYH2, MYH7* were associated with lower prognosis in HNSCC (head and neck squamous cell carcinoma (HNSCC), however, their expression is stage-dependent and is associated with immune infiltration as well [185, 186]. The role of *MYL11* (Myosin light chain 11) is not well established in the context of cancer. *MYL2* (Myosin light chain 2), a gene that is reported to be downregulated in rhabdomyosarcoma and associated with lower prognosis in these patients was also downregulated in this study, supporting its role as a TSG in sarcomas [187]. *MYH7* (Myosin Heavy Chain 7) gene has a negative correlation with lung cancer prognosis, nevertheless, its expression is again dependent on the stage of cancer [188]. Noteworthily, downregulated *MYL1* (Myosin light chain 1/3, skeletal muscle isoform), gene promotes tumor metastasis via tumor immune infiltration in HNSCC and its upregulation is associated with poor prognosis in HNSCC, and specifically in rhabdomyosarcoma (subtype of sarcoma more common among children) [189, 190]. It is evident that myosins have a diverse role in cancer regulating the tumor microenviroment in multiple ways and more studies are required to establish their role in sarcoma [191].

The expression of *MB* (Myoglobin) gene is established in sarcoma and it acts as a marker for ‘myoid differentiation’ in tumors of different origins [192]. Its loss in epithelial cancers such as breast and prostate cancer promotes partial mesenchymal transition from epithelial state resulting in migration [193, 194], and its high expression in breast cancer cells reduces cell migration through MB-driven oxidant signaling [195] establishing its TSG role. Additionally, reduced expression of *MB* is associated with poor prognosis in breast and prostate cancer [196, 197]. This downregulated gene could play a similar role in OA sarcoma patients. Interleukin-8 (*CXCL8*), an upregulated gene in OA sarcoma patients, is a chemotactic factor that mediates inflammatory response by attracting the neutrophils, basophils, and T-cells to clear the pathogens and protect the host from infection. Its overexpression plays a critical role in promoting tumorigenesis via tumor immunosuppression, and activation of tyrosine kinase pathways[198, 199]. High expression of this gene is associated with lower prognosis in liver, lung, and colorectal cancer [200–202].

Intriguingly, some of the hub genes like *ACTN2, TNNI2* and *TTN*, which are established to have a tumor promoting role in cancer is found to be downregulated in OA sarcoma patients. *ACTN2* has a pro-metastatic role in hepatocellular carcinoma (HCC) especially in later metastasis stages (extravasation and colonization) and also promotes cellular motility and invasion abilities in the same [203]. Troponin I, fast skeletal muscle (*TNNI2*), is the inhibitory subunit of troponin, conferring calcium sensitivity to striated muscle actomyosin ATPase activity. Its overexpression is associated with peritoneal metastasis and recurrence, and hence, it is proposed as a specific biomarker for gastric cancer [204]. Furthermore, it is linked with cell proliferation, migration, and invasion in pancreatic cancer cells and gastric cancer [205]. Titin (*TTN*) plays a role in the assembly and functioning of vertebrate striated muscles. It acts as a potential oncogene in CRC by promoting proliferation, invasion, and metastasis [206]. Further, *TTN* mutations were shown to be associated with immunotherapy response in multiple cancers [207, 208].

The identified hub genes are associated with tumor microenviroment via the regulation (tumor promotion or inhibition) of immune infiltration, EMT, and metastasis pathways. Hence, it is important to conduct more extensive studies that involve multi-omic analyses to understand the systemic role of these genes and validate the findings.

## 5 Conclusion and Future Perspectives

Sarcoma, a rare malignancy arising in the mesenchymal tissues of the body, is mainly subdivided into STS (soft tissue sarcoma) and BS (bone sarcoma). Excluding gastrointestinal sarcomas (having a net overall survival rate of 70% due to the development of targeted therapies), the survival rate of STS and BS has been stagnant for the past few years at 53% and 61% respectively [7]. Age has been reported to be a significant prognostic marker in many cancers and especially in sarcoma [6, 8, 13, 15–17, 43–45]. Our preliminary analysis, on par with some previous studies, showed a lower prognosis in OA (≥ 65 yrs) compared to YA patients (18-65 yrs) [7, 8, 13, 21, 22, 24, 25]. The lower prognosis of older age cancer patients, though mainly reported to be associated with undertreatment [18, 36–39], is also attributed to immunosenescence (decrease in anti-tumor cell-mediated immunity), alterations in ECM, increased inflammation observed in them [15, 46, 47]. Some studies reported an increased metastases potential in elderly cancer patients via remodeled ECM along with immunosenescence [48–50]. In the current study, it is observed that the significant dysregulated genes in the OA sarcoma patients were also associated with these pathways.

It is observed that many of the identified top 10 dysregulated (up/down-regulated) genes (*LINC02070, LINC01838, LOC105372937, ENSG00000286056, ENSG00000234803 (FAM197Y2), ENSG00000225560(FAM197Y8), ENSG00000286941, MIR202HG, FAM197Y7, OR4D10, OR7D4, ENSG00000286564, ENSG00000248958*) were novel findings including lncRNA’s, and not previously studied in the context of cancer. The previously reported oncogene *TRPM1* promoting tumor progression via calcium signaling in melanoma, was observed to be upregulated in OA sarcoma patients. The top 10 downregulated genes included *PGC, LIPF, SOX10, CACNG3, NCR2,* and *GKN2*, having a tumor suppressive role in other cancer types via regulating proliferation, metastases, EMT process, ECM degradation, and immune regulation [80–82, 89, 92–94]. Noteworthily, *PRB2* established to have a tumor suppressive role in cancer was upregulated in OA sarcoma patients, while *CRH* and the less studied *PMP2*, which are reported oncogenes, driving immune escape, angiogenesis and invasion in cancer, was found to be downregulated in the OA sarcoma patients [83, 84, 99]. These results indicate a possible cross-talk of multiple genes in the significant pathways controlling the tumor microenvironment. Hence, future studies that utilize multi-omics data of a larger cohort can shed more light on the role of these genes in a holistic view. Functional enrichment analysis of the sig-DEGs in OA patients revealed dysregulation of genes involved in MMP, ECM degradation, and EMT pathways, thus, associated with tumor microenviroment.

The 10 sig-TFs identified in this study were mostly downregulated (*CDX2, ELF3, GRHL2, HNF4A, OXT2, SOX10, SPDEF, PHOX2B*, and *PGR*) with only *NR5A1* upregulated in the OA patients. Some of them are studied previously in context to sarcomas such as *GRHL2, ELF3, SOX10, PGR* and *CDX2* [113–126]. It was observed that the majority of the downregulated TF’s (*GRHL2, ELF3, PGR, CDX2, HNF4A, SPDEF,* and *PHOX2B*) are potential TSGs in OA sarcoma patients, due to supporting evidence from previous studies either in sarcoma or other cancer types (Table 4.3.1). However, the upregulated *NR5A1* could be a potent oncogene in the same, due to its role in promoting self-renewing properties in stem cells, however needs further validation [112]. Interestingly, one of the sig-TFs *OTX2*, which is not studied in sarcoma, has an oncogenic role by promoting metastases in medulloblastomas [127, 128] was downregulated in OA sarcoma patients. Functional enrichment analysis and existing studies show that these sig-TF’s (*GRHL2, ELF3, SOX10, HNF4A, NR5A1* and *CDX2*) are associated with the promoting epithelial characteristics of the cells and inhibition of EMT, metastases and Wnt/*β*-catenin pathways [92, 113, 115–117, 119–125, 129].

Potential prognostic biomarkers were identified by the gene specific survival analysis of the OA sarcoma patients. Though none of the prognostic biomarkers identified in this study were reported to be associated with sarcoma, previous studies have established that the prognostic significance of potential TSG’s *BPIFB1, A4GNT, RHEX, CPA3, CX3CR1,* and oncogene *PLA2G1B* in multiple other cancers [145–156]. Some contradicting results were also observed, overexpression of *PPP1R17, EPS8L3,* and *CMA1* was reported to be associated with lower prognosis, however downregulation of these genes was significantly associated with lower prognosis of the OA sarcoma patients [157–159]. Although, the latter two are reported as oncogenes in other cancer types, *PPP1R17* has a dual role in cancer, and hence the role of *PPP1R17* in OA sarcoma should be further studied [148, 158, 159, 162]. FEA of the prognostic biomarkers revealed that the sig-survival genes were associated with angiotensin maturation, extracellular matrix interactions, and immune-mediated pathways, which is supported by previous research, with *RHEX, CX3CR1* and *CMA1* being associated with immune regulation pathways, *PLA2G1B* and *CMA3* are associated with ECM and angiotensin maturation [150–152, 159, 161]. The downregulated TSG’s *PPP1R17, BPIFB1,* and *TACR2* were known to inhibit tumorigenesis via inhibition of migration, and *β*-catenin signaling pathways [148, 166, 173].

The network analysis identified *MYH2, MYH7, MYL1, MYL2, MYL11, ACTN2, TTN, TNNI2, MB* and *CXCL8* as the top 10 hub genes. Though high expression of myosin genes (downregulated in this study) except *MYL11*, were shown to be associated with lower survival in HNSCC, they had a low expression in advanced stages (I, II, III, IV) and *MYL2* genes downregulation was associated with lower survival in rhabdomyosarcoma [187]. Hence, their role could be stage and tissue specific, requiring more extensive research. The upregulated *CXCL8* gene has a tumor promoting role (pro- metastatic, tumor infiltration) [198, 199] and associated with lower prognosis in other cancer types [200–202], while the downregulated MB shows a tumor suppressive role by inhibiting metastases and associated with lower prognosis in breast cancer[195–197]. Notably, the previously reported oncogenes (*TTN, ACTN2, TNNI2*) in other cancer types promoting metastases, were observed to be downregulated in our analysis [203–206]. It is observed that the hub genes play a role in the tumorigenesis process via regulation of tumor microenviroment.

The RNA-seq analysis of the TCGA-SARC dataset revealed a possible dysregulation of tumorigenic pathways such as EMT, Wnt/*β*-catenin pathway, tumor infiltration, angiogenesis, MMP, and ECM degradation pathways in OA patients. Many of the downregulated genes were associated with maintaining epithelial characteristics and their loss promoted mesenchymal transition. Previous studies have also reported an altered tumor microenvironment via EMC remodeling and increased inflammation, which can drive metastases in older cancer patients supporting the results from the current study. The identified genes and pathways could help in understanding the biology of OA patients and serve as biomarkers to develop new treatment regimens and improve efficacy. Additionally, these inferences also shed light on the role of these genes associated with lower prognosis in promoting tumorigenic pathways. Noteworthily, based on the existing studies in different cancer types, though majority of the identified dysregulated genes drive tumorigenic pathways (mentioned above) in OA sarcoma patients, respective dysregulation of KRT77 (Up), EPS8L3 (Down), OXT8 (Down), MYL1 (Down), ACTN2 (Down), TNNI2 (Down), and TTN (Down) genes are previously reported to alleviate the same. Hence, it is important to perform more extensive studies exploring multi-omics data of larger cohorts of OA sarcoma patients to check for cross-links between the genes and if they have dual roles in these pathways. The current study is limited by the sample size, diversity, and additional categorization of sarcoma, which could influence the results of the analysis. Future studies can validate the role of these genes in the tumor microenvironment and cancer progression in OA-sarcoma patients. Additionally, multi-omics analysis of these patients can provide more insights into the biological background for their lower prognosis, which needs to be further validated in clinical settings.

## 6 Abbreviations

BP: Biological process
BS: Bone sarcoma
CC: Cellular component
CCS: Clear-cell sarcoma
CRC: Colorectal cancer
EMT: Epithelial-to-mesenchymal transition
ECM: Extracellular matrix
FDR: False discovery rate
FEA: Functional enrichment analysis
DEG: Differential expressed genes
DGEA: Differential gene expression analysis
GO: Gene ontology
GPCR: G protein-coupled receptor
HCC: Hepatocellular carcinoma
HNSCC: Head and neck squamous cell carcinoma
HRs: Hazard ratios
KEGG: Kyoto Encyclopedia of Genes and Genomes
KM: Kaplan-Meier
lncRNA: Long non-coding RNA
logFC: Log fold-change
MCC: Maximal Clique Centrality
MF: Molecular function
MMP: Matrix metalloproteinases
MPNST: Malignant peripheral nerve sheath tumor
OA: Older age patients (above the age of 65)
OS: Overall survival
PPI: Protein-protein interactions
RNA: Ribonucleic acid
SARC: Sarcoma (in TCGA database)
Sig-DEGs: Significant differentially expressed genes
Sig-TFs: Significant differentially regulated TFs
Sig-Survival: Significant differentially expressed genes associated with OA patient survival
STRING: Search Tool for the Retrieval of Interacting Genes/Proteins
STS: Soft-tissue sarcomas
TCGA: The Cancer Genome Atlas
TF: Transcription factor
TFEA: Transcription factor enrichment analysis
TG: Target gene
TME: Tumor microenvironment
TSG: Tumor suppressive gene
YA: Younger age patients (between 18-65 years)

## Acknowledgements

The authors would like to express their gratitude to Adewale Ogunleye and Richard Agyekum from the HackBio team for their mentorship and inspiring us to complete this research project.

## Author Contributions

Vidhyavathy Nagarajan: Conceptualized and designed the study; Developed the code, Analyzed, visualized and interpreted the data; wrote and reviewed the manuscript.

Shreya S. Karandikar and Mary S.J. Dhevanayagam: Analyzed and visualized the data; wrote and reviewed the manuscript.

## Data Availability and Code

The datasets supporting the findings of this study and the code used for the analysis is available on the Github repository.

