## Supplemental Figures and Table for "Transcriptomic Profiling of Old Age Sarcoma Patients using TCGA RNA-seq data"

### Supplementary Data

#### Supplementary Figures

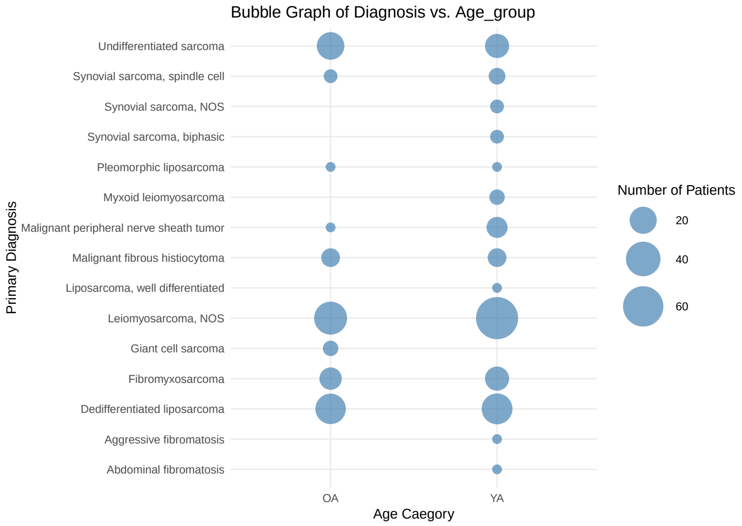


**Supplementary Figure 1.** Patient Demographics representing the distribution of subtypes among the OA and YA patient groups.

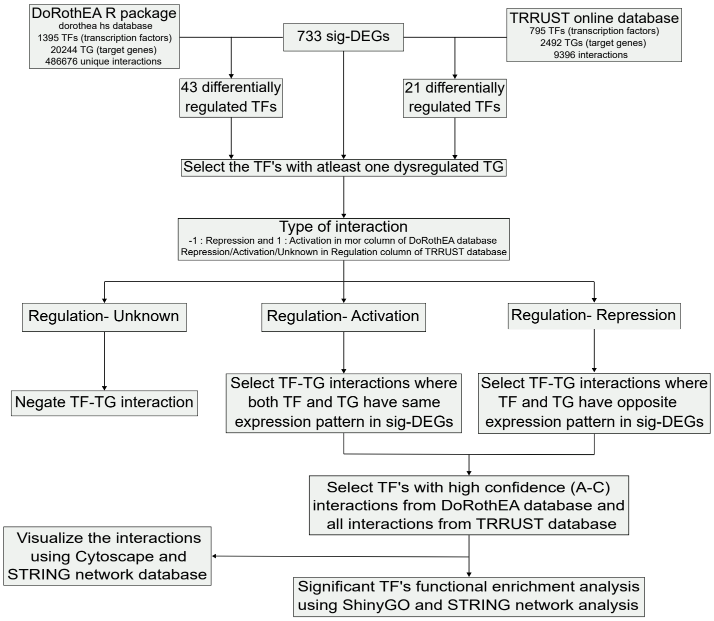


**Supplementary Figure 2.** Pipeline for Transcription factor enrichment analysis (TFEA) to identify significant transcription factors differentially expressed in OA patients.

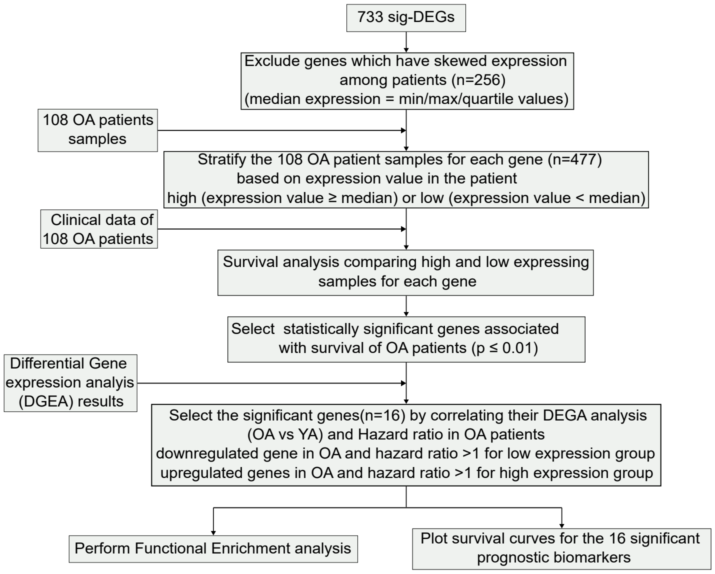


**Supplementary Figure 3.** Pipeline for Gene Specific Survival analysis to identify significant prognostic markers in OA patients.

### Supplementary Tables

| Top 10 in network STRING network - SigDEG ranked by Degree method | | | |
|---|---|---|---|
| Rank | Name | Gene | Score |
| 1 | 9606.ENSP00000347507 | MYH7 | 41 |
| 2 | 9606.ENSP00000325239 | MYL11 | 40 |
| 2 | 9606.ENSP00000307280 | MYL1 | 40 |
| 4 | 9606.ENSP00000467141 | TTN | 39 |
| 4 | 9606.ENSP00000355537 | ACTN2 | 39 |
| 4 | 9606.ENSP00000380367 | MYH2 | 39 |
| 7 | 9606.ENSP00000352835 | MB | 38 |
| 8 | 9606.ENSP00000371336 | TNNI2 | 37 |
| 8 | 9606.ENSP00000228841 | MYL2 | 37 |
| 8 | 9606.ENSP00000306512 | CXCL8 | 37 |
| **Top 10 in network STRING network - SigDEG ranked by Closeness method** | | | |
| Rank | Name | Gene | Score |
| 1 | 9606.ENSP00000306512 | CXCL8 | 193.6333333 |
| 2 | 9606.ENSP00000253408 | GFAP | 190.7 |
| 3 | 9606.ENSP00000297261 | SHH | 188.8833333 |
| 4 | 9606.ENSP00000321106 | TAC1 | 184.2 |
| 5 | 9606.ENSP00000216200 | PVALB | 183.0095238 |
| 6 | 9606.ENSP00000216492 | CHGA | 179.3428571 |
| 7 | 9606.ENSP00000362108 | LCN2 | 179.0333333 |
| 8 | 9606.ENSP00000263735 | EPCAM | 178.1 |
| 9 | 9606.ENSP00000356438 | PTGS2 | 177.9333333 |
| 10 | 9606.ENSP00000168712 | FGF4 | 177.8833333 |
| **Top 10 in network STRING network - SigDEG ranked by MCC method** | | | |
| Rank | Name | Gene | Score |
| 1 | 9606.ENSP00000355537 | ACTN2 | 4.50E+21 |
| 2 | 9606.ENSP00000467141 | TTN | 4.50E+21 |
| 3 | 9606.ENSP00000325239 | MYL11 | 4.50E+21 |
| 3 | 9606.ENSP00000307280 | MYL1 | 4.50E+21 |
| 5 | 9606.ENSP00000347507 | MYH7 | 4.50E+21 |
| 6 | 9606.ENSP00000228841 | MYL2 | 4.50E+21 |
| 7 | 9606.ENSP00000380367 | MYH2 | 4.50E+21 |
| 8 | 9606.ENSP00000484342 | NEB | 4.50E+21 |
| 9 | 9606.ENSP00000357057 | CASQ1 | 4.50E+21 |
| 10 | 9606.ENSP00000386041 | MYH6 | 4.50E+21 |

**Supplementary Table 1.** Rank of top 10 hub genes in each algorithm
